# Single-nucleotide substitution determines pollen production in Japanese cedar

**DOI:** 10.1101/2022.03.17.484665

**Authors:** Hiroyuki Kakui, Tokuko Ujino-Ihara, Yoichi Hasegawa, Eriko Tsurisaki, Norihiro Futamura, Junji Iwai, Yuumi Higuchi, Takeshi Fujino, Yutaka Suzuki, Masahiro Kasahara, Katsushi Yamaguchi, Shuji Shigenobu, Masahiro Otani, Masaru Nakano, Saneyoshi Ueno, Yoshinari Moriguchi

## Abstract

Pollinosis, also known as pollen allergy or hay fever, is a global problem caused by pollen produced by various plant species^1–6^. The wind-pollinated Japanese cedar (*Cryptomeria japonica*) is the largest contributor to severe pollinosis in Japan, where increasing proportions of people have been affected in recent decades^7^. The *MS4* (*MALE STERILITY 4*) locus of Japanese cedar controls pollen production, and its homozygous mutants (*ms4/ms4*) show abnormal pollen development after the tetrad stage and produce no mature pollen. In this study, we narrowed down the *MS4* locus by fine mapping in Japanese cedar and found *TKPR1* (*TETRAKETIDE α-PYRONE REDUCTASE*) gene in this region. Transformation experiments using *Arabidopsis thaliana* showed that single-nucleotide substitution of *CjTKPR1* determines pollen production. Broad conservation of TKPR1 beyond plant division could lead to the creation of pollen-free plant not only for Japanese cedar but also for broader plant species.

## Introduction

Pollen production is critical for plant reproduction; therefore, the genetics of pollen development have been studied extensively. This research has led to the discovery of the pollen developmental pathway, as well as many genes involved in pollen development regulation^8^. Several gene mutants give rise to pollen-free phenotype called male sterility^8,9^. In agriculture, pollen production is also important. Many pollen grains are required for artificial pollination and efficient seed/fruit production. Conversely, male sterility is a crucial trait required to produce hybrid crops with stable quality and quantity^10^. For our health, pollen can sometimes be harmful by triggering an allergic reaction, pollinosis^1,11^.

Pollinosis is a global problem caused by pollen from various plant species, including both angiosperms and gymnosperms^1,11^. For example, the prevalence of pollinosis is estimated at 40% in Europe^1^. The global pollen load has increased in recent decades that thought mainly driven by climate change^12,13^. Species of the family Cupressaceae, which belong to gymnosperms, are widely distributed over the eastern Mediterranean, Asia, and western North America. These species have long been used for timber or pharmaceutical products (e.g., essential oils, fragrances, and folk medicines); however, pollen produced by some Cupressaceae species are major triggers of severe pollinosis; these include Arizona cypress (*Cupressus arizonica*), Italian cypress (*Cupressus sempervierens*), mountain cedar (*Juniperus ashei*), Japanese cypress (*Chamaecyparis obtusa*), and Japanese cedar (*Cryptomeria japonica*)^6,14^.

Japanese cedar is a wind-pollinated conifer, and is the most important timber species in Japan, accounting for 58% (12.7 million m^3^) of total roundwood production in 2019^15^. However, Japanese cedar is also the leading cause of the most severe allergic reactions to pollen in Japan^7^. A single Japanese cedar tree typically produces thousands of male strobili, which correspond to male flowers in angiosperms; a single male strobilus contains approximately 100,000–300,000 pollen grains (Fig. 1a–c, Supplementary Video 1)^16^. Upon release, these vast numbers of pollen grains are carried away by the wind. The proportion of people suffering from pollinosis caused by Japanese cedar pollen in Japan has increased in recent years, from 16.2% in 1998 to 26.5% in 2008, and 38.8% in 2019^17,18^. Therefore, planting pollen-free trees is a promising strategy for maintaining Japanese cedar as a building material while countering its effects in pollinosis.

**Fig. 1.**
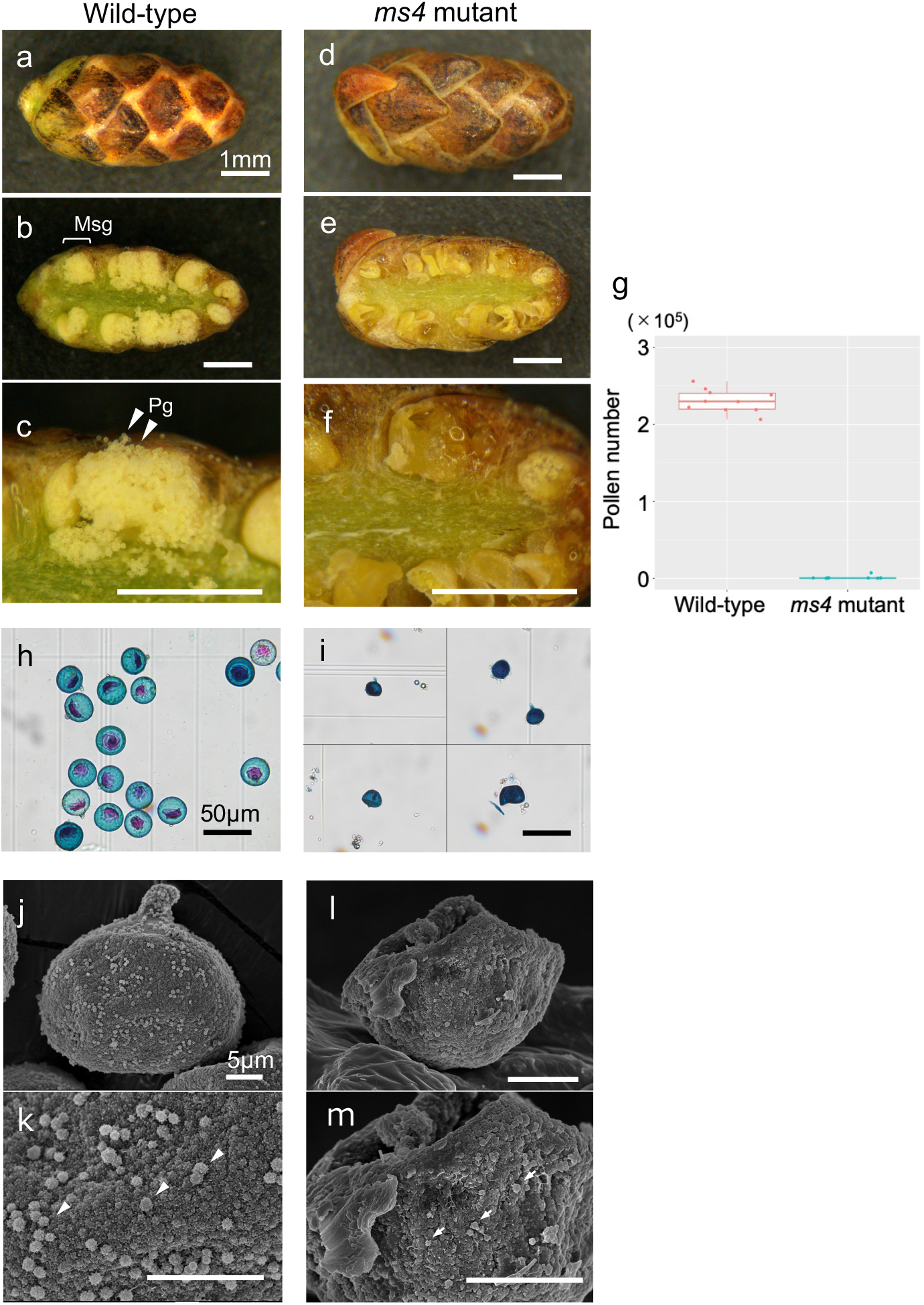
Male strobili and pollen from wild-type (WT) and *ms4* mutant Japanese cedar plants. Male strobili of the WT (‘Higashikanbara-5’, **a–c**) and *ms4* mutant (‘Shindai-8’, **d–f**). Scale bars, 1 mm. Transverse-cut sample from the WT (b, c) showed many pollen grains (Pg) in each microsporangium (Msg), whereas the *ms4* mutant had no or few pollen grains in microsporangia (**e, f**). **g**, Pollen grain count for a single male strobilus. Although approximately 200,000 pollen grains were produced in the WT, few pollen grains (median = 0–7,000) were counted in the *ms4* mutant (*n* = 10, 10). **h, i**, Pollen grain observations following Alexander staining. WT plants produced globular pollen grains with single projections (papillae) (**h**), whereas *ms4* plants produced small, squashed pollen grains. Scale bars, 50 μm. **j–m**, Scanning electron microscopy (SEM) observations of normal (**j, k**) and *ms4* mutant (**l, m**) pollen. Although normal pollen had uniform orbicules (arrows, **k**), *ms4* mutant pollen had small, irregular orbicules (arrows, **m**). Scale bars, 5 μm. Data shown in **g** are provided in a source data file.

Conifers have large genomes, which complicates genetic studies of these species (e.g., 25.4 Gb for *Pinus tabuliformis*^19^ and ∼20 Gb for *Pinus taeda*^20^). Flowering trait analyses are particularly problematic due to the necessity of large fields for keeping materials for segregation analyses, as well as a wait time of several years until the flowering stage can be examined^21^. Among conifers, Japanese cedar has several advantages for flowering trait analyses. Its genome is smaller (∼10.8 Gb)^22^ than those of other conifer species, and its flowering can be induced artificially using gibberellic acid^23^. Previous field studies identified four male sterility loci, i.e., *MALE STERILITY 1–4* (*MS1–MS4*), in Japanese cedar^24,25^, through linkage analyses^25,26^. Recently, a strong candidate gene for *MS1* was discovered by our research^27^. Two different mutation alleles of *CJt020762*, which is located on the *MS1* locus, showed pollen-free phenotype, suggesting that *CJt020762* is a causal gene of *MS1*, although functional validation is required.

## Results

The *ms4* mutant ‘Shindai-8’ was discovered in a planted forest in Niigata Prefecture, Japan. Mature male strobili of homozygous *ms4* mutants (*ms4/ms4*) produced almost no pollen (Fig. 1a–g). Very little of the pollen produced by the *ms4* mutants showed obvious abnormal phenotypes compared to wild-type (WT) plants, such as small or collapsed pollen grains (Fig. 1h, i). Scanning electron microscopy (SEM) observations revealed that the *ms4* mutant produces small, abnormal orbicules, which are components of sporopollenin (Fig. 1j–m)^28^. Sporopollenin is an essential constituent of the pollen wall, and dysfunctional sporopollenin biosynthesis genes often result in male sterility^29,30^.

To narrow down the region of *MS4* locus, we generated a segregating population by crossing the *ms4* mutant with heterozygous individuals (‘Sindai-8’ × ‘S8HK5’, Supplementary Table 1). We found that 46 individuals produced normal pollen, whereas 48 showed pollen-free phenotype (Supplementary Table 2). Then, we conducted fine mapping using these 94 individuals and narrowed down *MS4* for approximately 7.65 Mb-wide region on linkage group 4 (Fig. 2a). Annotation analysis revealed 67 genes in this region (Fig. 2b, Supplementary Table 3). Among these genes, we found a homologous gene of *TKPR1* (*TETRAKETIDE α-PYRONE REDUCTASE 1* (*TKPR1*, hereafter *CjTKPR1*); this gene is essential for sporopollenin biosynthesis through making the exine layer of the pollen wall^29,31^. RNA sequencing (RNA-seq) data revealed that *CjTKPR1* had the highest expression levels in all three male strobilus samples (183.17–1131.88 transcripts per million (TPM)), but less or no expression in inner bark and leaf tissues (0 and 0.54 TPM, respectively; Fig. 2b and Supplementary Table 3)^32^.

**Fig. 2.**
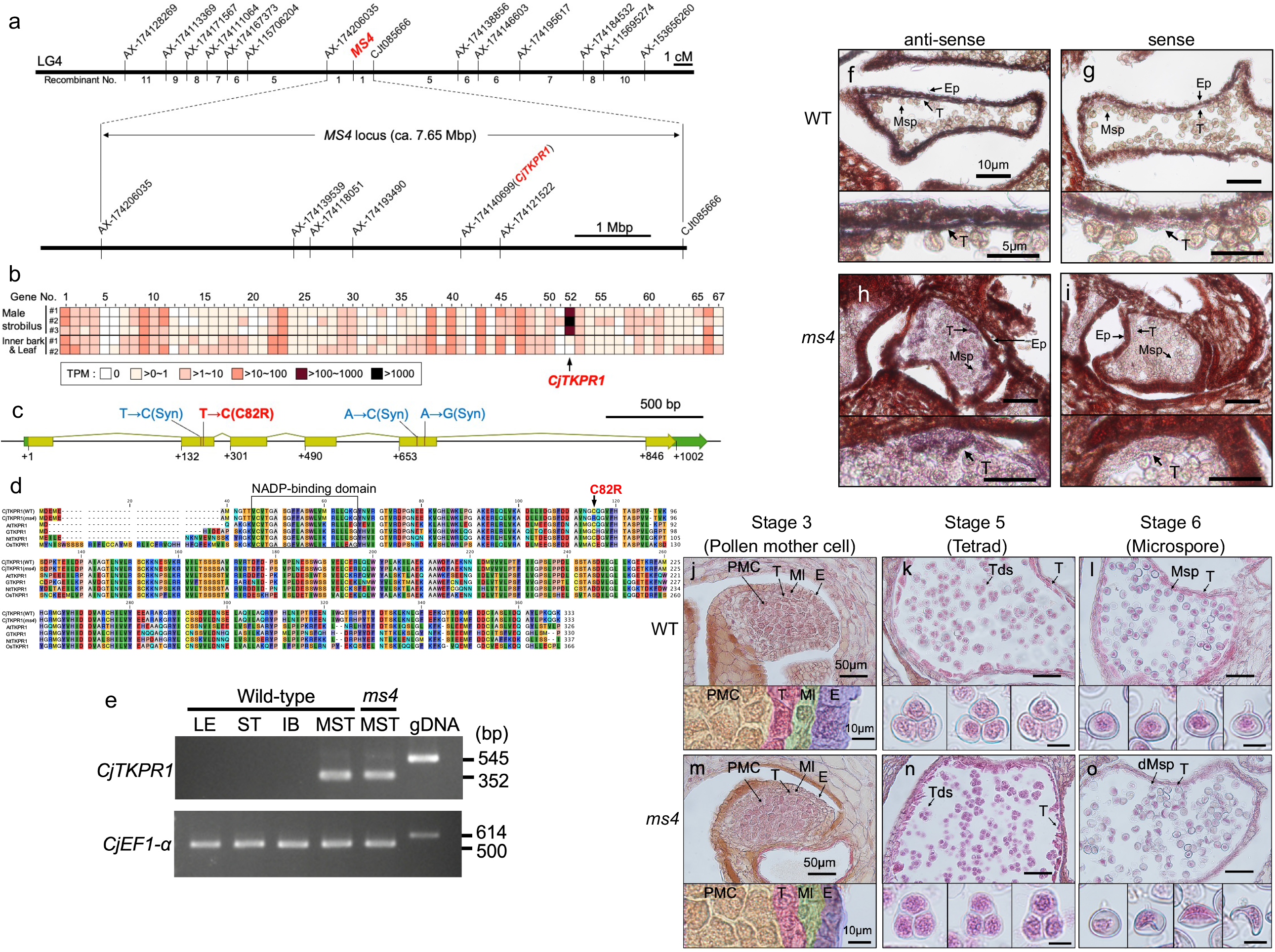
Fine mapping and analysis of candidate genes in the *MS4* locus. **a**, Fine mapping of the *MS4* locus according to a linkage map around *MS4* (top) and physical map of the *MS4* region (bottom). **b**, Expression patterns of 67 genes of the *MS4* region from male strobili (three samples) or inner bark and leaf tissues (two samples). RNA sequencing (RNA-seq) data were obtained from our previous study^32^ and expression levels were categorised into six levels (0, > 0–1, > 1–10, > 10–100, > 100–1,000, or > 1,000 transcripts per million, TPM). Gene no. 52, *CjTKPR1*, had the highest expression level among male strobilus samples but no or little expression was detected in inner bark or leaf tissues. Precise expression data for each gene are provided in Supplementary Table 3. **c**, Gene structure of *CjTKPR1*. Green and yellow boxes indicate mRNA and coding regions, respectively. Four nucleotide substitutions between of *ms4* mutant (‘Shindai-8’) are shown above the boxes. Only a single amino acid substitution was detected at position 244 nt (C82R); the others were synonymous mutations (Syn). **d**, Amino acid comparison among CjTKPR1, AtTKPR1, GTKPR1, NtTKPR1, and OsTKPR1. A single amino acid substitution of *ms4* mutant (C82R, arrow) was conserved within various plant species (Supplementary Fig. 1). The putative NADP-binding domain is framed. Multiple-sequence alignment was generated using CLC Main Workbench v22.0 software. **e**, Reverse-transcription polymerase chain reaction (RT-PCR) of the *CjTKPR1* transcript. *ELONGATION FACTOR1-alpha* (*CjEF1-α*) was used as an internal control. gDNA, genomic DNA; IB, inner bark; LE, leaf; MST, male strobilus; ST, stem. **f–i**, *in situ* hybridisation of *CjTKPR1*. Magnified views are shown at the bottom of each figure. *CjTKPR1* expression was observed in the tapetum (T) of both WT (**f**) and *ms4* mutant plants (**g**), whereas no signal was observed from the sense probe (**g, i**). Scale bars: top = 10 μm, bottom = 5 μm. **j–o**, Transverse section of male strobilus development in WT (**j–l**) and *ms4* mutant plants (**m–o**). Microsporangia of male strobili from stages 3 (**j, m**) 5 (**k, n**), and 6 (**l, o**). Magnified views are shown at the bottom of each figure. Male strobilus stages were defined previously^45^. dMsp, degraded microspores; E, epidermis; Ml, middle layer; Msp, microspore; PMC, pollen mother cell; T, tapetum; Tds, tetrads. Scale bars: top = 50 μm, bottom = 10 μm. Data shown in **a** are provided in a source data file.

*TKPR1* is expressed in the tapetum, and tetraketide reductase activity of TKPR1 was observed *in vitro* for *Arabidopsis thaliana* and tobacco^33,34^. Mutants of *TKPR1* lead to male sterility in several angiosperm species such as *A. thaliana*, rice, tobacco, and gerbera^31,33–35^. Histological observations have revealed that mutants of *AtTKPR1* and *OsTKPR1* show no obvious defects at the tetrad stage, although abnormal microspores appeared in later stages and pollen cells, before degenerating rapidly^35,36^. CjTKPR1 shares 54.6–64.9% amino acid identity with *A. thaliana*, gerbera, tobacco, and rice (Fig. 2c, d, and Supplementary Fig. 1). Reverse-transcription polymerase chain reaction (RT-PCR) showed specific *CjTKPR1* expression in the male strobilus from wild-type and *ms4* mutant (Fig. 2e). Furthermore, *in situ* hybridisation of male strobili confirmed *CjTKPR1* expression in the tapetum of Japanese cedar (Fig. 2f–i). Histological observations of male strobili showed that the *ms4* mutant has a clear tetrad structure, but also has crushed, abnormal microspores (Fig. 2j–o) as previously observed in *A. thaliana* and rice^35,36^. These results support the hypothesis that *CjTKPR1* is the causal gene of *MS4* in Japanese cedar.

Sequence comparison of *CjTKPR1* between WT (‘Higashikanbara-5’) and *ms4* (‘Shindai-8’) revealed only four nucleic acid substitutions; interestingly, only one amino acid substitution was detected at position 244 nt (C82R, Fig. 2c, d) among these substitutions. This single-nucleotide polymorphism (SNP) at 244 nt was completely linked with the pollen-free phenotype in wild accessions (Supplementary Table 1) and two crossing progenies (‘S8’ × ‘S8HK5’, Supplementary Table 2; selfing progenies of ‘S8DY1’, Supplementary Table 4), except for mutants from other male sterility loci (*ms1–ms3*). Thus, only plants with C/C at position 244 nt showed the pollen-free phenotype, and those having T/T or T/C showed normal pollen production (Supplementary Table 1, 2, and 4). Furthermore the cysteine-82 of CjTKPR1 was perfectly conserved in various plant species, including both angiosperms and gymnosperms (Fig. 2d and Supplementary Fig. 1). These results suggest that *CjTKPR1* is an important gene for pollen production in Japanese cedar, and that the nucleotide substitution at position 244 nt (C82R) is a strong candidate for the causal mutation of *MS4* in Japanese cedar.

Therefore, we conducted complementation tests to confirm that *CjTKPR1* sequences control pollen production using *A. thaliana* (Fig. 3). First, we obtained the *TKPR1* mutant of *A. thaliana* (*Attkpr1-1* mutant, SAIL_837_D01), which homozygous plant shows a pollen-free phenotype^33^ (Fig. 3b, c). Then, four *CjTKPR1* sequences, i.e., the WT, *ms4* mutant (MT), WT background with a point mutation at 244 nt (T to C, WM), and *ms4* background with a point mutation at 244 nt (C to T, MW) (Fig. 3a), were each introduced into heterozygous plants of *AtTKPR1 (AtTKPR1/Attkpr1-1)*, not into *Attkpr1-1* homozygous plants *(Attkpr1-1/Attkpr1-1)* that produce no pollen and therefore obtain no progenies. Next, we selected *Attkpr1-1* homozygous plants with *CjTKPR1* constructs from the T2 or T3 generations. Anthers of these plants were stained by Alexander staining and pollen production was confirmed^37^. As a result, *CjTKPR1-WT*-introduced plants recovered pollen production (Fig. 3d), whereas *CjTKPR1-MT*-introduced plants still showed the pollen-free phenotype (Fig. 3e). These results indicate that *CjTKPR1-WT* can complement pollen production in *A. thaliana*, and that the *CjTKPR1* allelic difference controls pollen production. Lines with a single-nucleotide substitution at 244 nt showed that *CjTKPR1-WM*-introduced plants, which have a *CjTKPR1-WT* background with a point mutation at 244 nt, could not recover pollen production (Fig. 3f). However, *CjTKPR1-MW*-introduced plants, which have a *ms4* background with a point mutation at 244 nt, recovered pollen production (Fig. 3g). Together, these complementation test results indicate that single-nucleotide substitution of *CjTKPR1* at 244 nt determines pollen production.

**Fig. 3.**
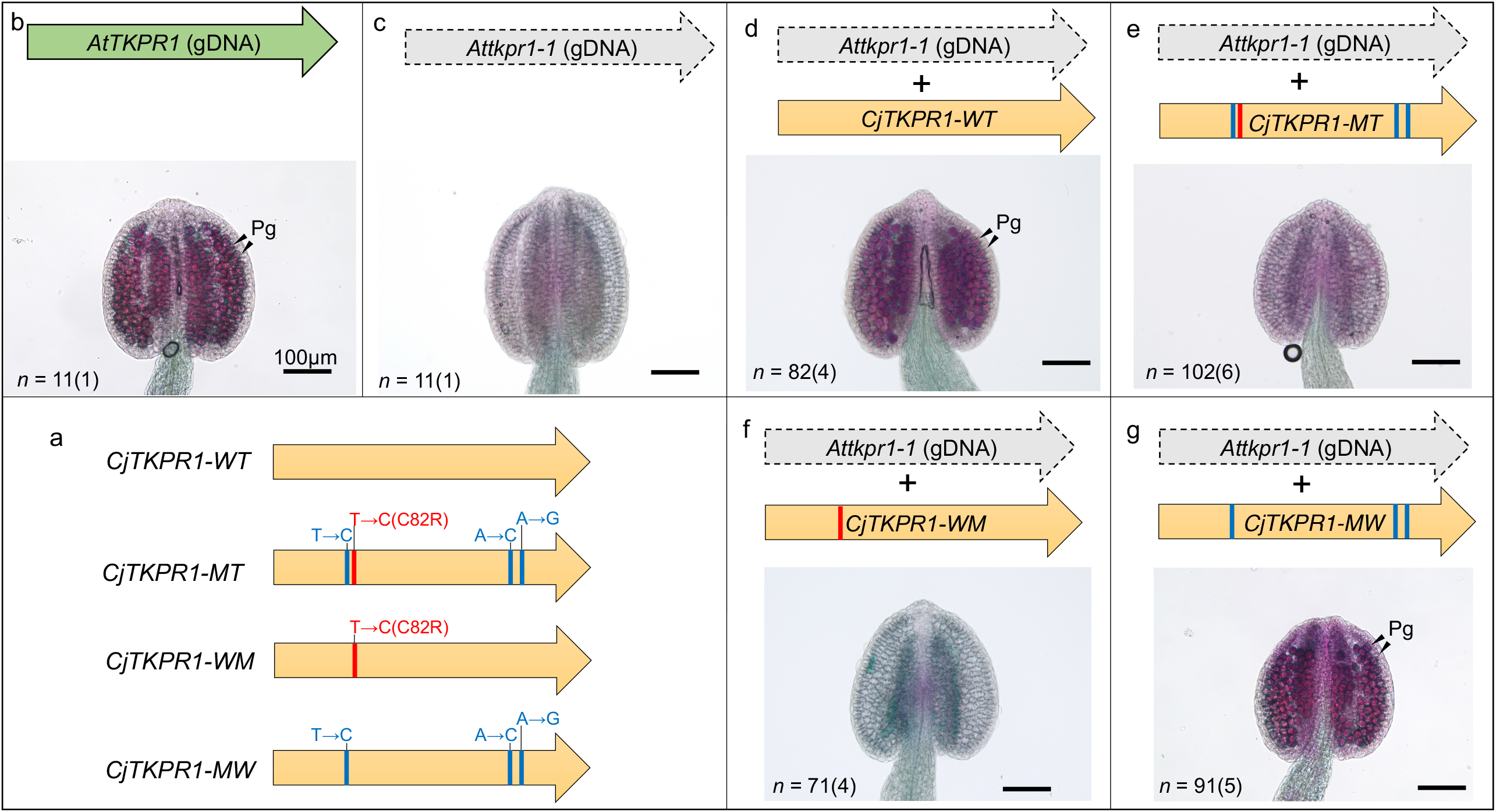
Complementation analysis of *CjTKPR1*. Four types of *CjTKPR1* sequences were constructed (**a**) and transgene mutants were prepared in the *Attkpr1-1/ Attkpr1-1* background. Red and blue lines indicate *ms4* single-nucleotide polymorphisms (SNPs), where red indicates a single-amino acid substitution (C82R, 244 nt position). The *Attkpr1-1* homozygous mutant showed pollen-free phenotype in contrast to the WT (**b, c**). Plants inserted with *CjTKPR1-WT* and *CjTKPR1-MW* (with T at 244 nt) recovered their pollen production (**d, g**), whereas those inserted with *CjTKPR1-MT* and *CjTKPR1-WM* (with C at 244 nt) showed no pollen production (**e, f**). Expressions of the inserted *CjTKPR1* were confirmed by RT-PCR (Supplementary Fig. 2). Numbers indicate anther counts, followed by the number of biological replicates in parentheses. Data shown in **b–g** are provided in a source data file. Pg: Pollen grain.

In this study, we identified a single-nucleotide substitution of *CjTKPR1* as a causal mutation of *MS4* in Japanese cedar. This knowledge provides the basic information for selecting and breeding pollen-free Japanese cedar efficiently and precisely. Our results also contribute for the creating new pollen-free plant using genome editing by targeting *TKPR1* gene. Cupressaceae species are suitable target plant species, which are often used for timber but are also global sources of severe pollinosis, include Arizona cypress, Italian cypress, and mountain cedar^14^. Since the recent progress of genome-editing methods that enable the application of mutagenesis for broader plant species, including Japanese cedar^38^, combining our results with genome editing methods will accelerate the creation of new pollen-free plants.

Tapetum expression of *CjTKPR1* in both WT and *ms4* mutant plants suggests that *CjTKPR1* of *ms4* may be dysfunctional at the protein level, but not at the transcription level, for example in terms of tetraketide reductase activity (Fig. 2e–i). This is the first study to identify dysfunction of TKPR1 activity at the single amino acid level. Because the activity domain of TKPR1 has not yet been analysed well, our results give a new insight into the enzymatically important amino residues for tetraketide reductase activity of TKPR1.

Interestingly, TKPR1 occurs not only in seed plants, but also in phylogenetically basal plants without pollen, such as the fern *Ceratopteris richardii* and liverwort *Marchantia polymorpha* (Supplementary Fig. 1). Although ferns and liverworts produce sperm as male gametes and do not have a pollen wall structure, they have a perispore layer within the spore that consists of sporopollenin. Sporopollenin is thought to be a key structure for the colonisation of terrestrial environments during plant evolution, protecting them from ultraviolet B light^39,40^. Our results that *CjTKPR1* could rescue disfunction of *TKPR1* of *Arabidopsis thaliana*, may imply other pollen-developmental genes are also conserved among the Plantae. Further investigation of the pollen developmental genes of Japanese cedar and comparison thereof among other plant species will help elucidate the evolution of male gametes in plants.

## Methods

### Plant Materials

Male strobili of Japanese cedar were collected from planted fields. We used 10 lines of wild accessions, 3 crossing parents (Supplementary Table 1), and two combinations of crossing progenies (Supplementary Table 2 and 4). Detailed information including sample name, sampling location, pollen phenotype, and SNP types at 244 nt of *CjTKPR1* for all the Japanese cedar materials used in this study are listed in Supplementary Table 1, 2, and 4. DNA and RNA were extracted from leaf^26^ and male strobilus^27^ tissues, as described previously.

### Male strobilus and pollen observations

For the male strobilus observations, WT (‘Higashikanbara-5’) and *ms4* mutant (‘Shindai-8’) of Japanese cedar plants were used. Male strobili were cut using a razor and photographed under a stereomicroscope (SZ-11; Olympus, Tokyo, Japan) using a digital camera (Wraycam EL310; Wraymer, Osaka, Japan). Pollen grains were stained with Alexander staining solution^37^, and photographed under a light microscope (BX-50; Olympus). Pollen grains from ‘Higashikanbara-5’ and ‘Shindai-8’ were counted using a cell counter^41^. Briefly, a single male strobilus was gently squashed using a pestle and suspended with water. The pollen suspension was then filtrated through two mesh filters (100 and 20 μm) to remove male strobilus debris and dust. The cleaned suspension was mixed with a cell counter buffer (CASYton; OLS, Bremen, Germany), and pollen grains were counted using a CASY cell counter (OLS).

### SEM observation

Pollen grains were extracted from progenies of ‘S8’ × ‘S8HK5’ (‘P411’ as normal pollen and ‘P380’ as *ms4* mutant, Supplementary Table 1). Male strobili were harvested on 12 December, 2019, after pollen maturation but before pollen release. After washing with deionised water, the male strobili were air-dried for 2 days. Pollen was harvested from dried male strobili and further sifted through a 75-μm mesh. The samples were coated using an osmium plasma ion coater (OPC80N; Filgen, Nagoya, Japan) and SEM observations were performed using a JSM-7500F system (JEOL, Tokyo, Japan) at 5 kV.

### Fine mapping of the *MS4* locus

Linkage maps were constructed using a *MS4* mapping family consisting of 46 normal pollen production offspring and 48 male sterile offspring produced by artificial crossing (‘S8’ × ‘S8HK5’; Supplementary Table 2). The SNP marker AX-174140699 on *CjTKPR1* was previously mapped on LG4 with five adjacent markers^26^. These markers were developed from transcript sequences blasted against two versions (0.11 and 0.1) of draft genome assemblies of *C. japonica*, leading to the identification of three genomic contig sequences (ctg370 and ctg2011 from ver. 0.11, and ctg482 from ver. 0.1). These three genomic contig sequences were manually combined and genomic coordinates were calculated, starting from AX-174206035 (Fig. 2a, Supplementary Table 3). Transcript sequences from *C. japonica* (CJ3006NRE)^32^ were mapped against genomic contig sequences using the minimap2 tool and the “-ax splice” option, resulting in 67 transcripts (Supplementary Table 3). An additional marker (CJt085666, Supplementary Table 3) was genotyped for the mapping family using PCR–restriction fragment length polymorphism (RFLP) and the *Nde*I restriction enzyme, to obtain *CjTKPR1* and the genetic markers on either side (Fig. 2a).

### RT-PCR

cDNA was synthesised from 500-ng aliquots of total RNA from Japanese cedar and *A. thaliana* using the PrimeScript IV 1^st^ Strand cDNA synthesis kit (Takara Bio, Shiga, Japan). *CjTKPR1* and *EF1-α* were amplified using intron-containing primers to distinguish between cDNA and gDNA amplification. The primer sequences are listed in Supplementary Table 5.

### *in situ* hybridisation

Full-length cDNA of *CjTKPR1* was cloned into the pZErO vector (Invitrogen, Waltham, MA, USA). Subsequently, 150-bp DNA fragments were amplified from the cloned full-length cDNA using gene-specific primers containing SP6 or T7 RNA polymerase promoter sequences at the 5’-end (Supplementary Table 5). PCR products were purified through phenol/chloroform/isoamyl alcohol extraction followed by ethanol precipitation, and used as templates for *in vitro* transcription. Sense and anti-sense digoxigenin-labelled probes were synthesised using the DIG RNA Labeling Kit (SP6/T7; Roche, Basel, Switzerland) according to the manufacturer’s instructions.

Male strobili of WT (‘Nakakubiki-4’) and *ms4* mutant (‘P380’) plants were fixed in FAA solution (formaldehyde:acetic acid:70% ethanol = 5:5:90) at 4°C for 24 h. After dehydration and embedding, the strobili were cut to a thickness of 20 μm. Hybridisation and immunological detection were performed as described previously^42,43^.

### Histological analysis of male strobili

Male strobili were collected from ‘Santo-1’ (WT) and ‘Shindai-8’ (*ms4* mutant) plants, which were grown at the Niigata Prefectural Forest Research Institute. Male strobili were fixed with FAA, dehydrated using graded ethanol series, introduced to chloroform through a chloroform–ethanol series, and embedded in paraffin. We cut 10-μm sections using a microtome (ESM-100L; ERMA, Tokyo, Japan), which were then stained with Mayer’s hematoxylin solution and mounted in Eukitt mounting medium (ORSAtec, Bobingen, Germany). These sections were observed under a light microscope (BX-50; Olympus) equipped with a digital camera (Wraycam-NOA2000; Wraymer).

### Transgenic experiments

Total RNA of *CjTKPR1* was extracted from male strobili of *Ms4/ms4* heterozygous individual (‘S8TM4’, Supplementary Table 1). cDNA was synthesised using the PrimeScript II kit (Takara Bio). Total RNA and genomic DNA from *A. thaliana* (Col-0 accession) were extracted from flower bud and leaf tissues, respectively. Coding regions of *CjTKPR1* and *AtTKPR1* were amplified using PrimeStar GXL polymerase (Takara Bio) with specific primers (Supplementary Table 5). pENTR1a, *AtTKPR1* promoter (pro), coding sequences of *CjTKPR1* or *AtTKPR1*, and Nos terminator (NosT) fragments were combined with In-Fusion HD mix (Takara Bio) and cloned to *Escherichia coli*. Single-nucleotide substitution constructs were made by inverse PCR with mutation-containing primers (Supplementary Table 5). Each AtTKPR1pro:TKPR1 coding sequence:NosT construct was further cloned into a pFAST-G01 vector using Gateway LR clonase II (Thermo Fisher Scientific, Waltham, MA, USA). We prepared *AtTKPR1/Attkpr1-1* heterozygous plants (Col-0 background, accession no. CS837358) as the transformation hosts because the *Attkpr1-1* homozygous mutant produces no pollen and therefore no progenies. Each construct was transformed into *AtTKPR1* heterozygous mutants via *Agrobacterium tumefaciens* (GV3101) using the floral dip method^44^. *AtTKPR1* heterozygous individuals containing an external *TKPR1* construct were selected based on fluorescence in the seed and confirmed by PCR (Supplementary Table 5). For the next generation, we selected *Attkpr1-1* homozygous mutants with external *TKPR1* sequences. Pollen production was analysed based on anther observation under a microscope after Alexander staining^37^.

## Supplementary Information

Supplementary Figures (S1, S2)

Supplementary Tables (S1-S5)

Supplementary Video 1

## Acknowledgements

We thank Shuh-ichi Nishikawa, Tsuyoshi Nakagawa, Yamagata Prefectural Forest Research and Instruction Center, and Ishikawa Agriculture and Forestry Research Center Forestry Experiment Station for providing materials, Momoko Ikeuchi, Toshiaki Tameshige, for technical support, Daisuke Maruyama, and Naoto-Benjamin Hamaya for valuable suggestions. Computation time was provided in part by the Data Integration and Analysis Facility, National Institute for Basic Biology, and by the SuperComputer System, and by Institute for Chemical Research, Kyoto University. This study was supported by JSPS KAKENHI (19K05976 to HK, 20H03239 to MK), FFPRI Grant (#201421, #201906) and NIBB Collaborative Research Program (16-403, 17-405, 18-408) to SU, Bio-oriented Technology Research Advancement Institution (BRAIN) Grant (JPJ007097; Project ID 28013B) to YM.

## Author contributions

HK, TUI, SU, and YM conceived and designed the study; HK, TUI, YHa, NF, JI, YHi, SU, and YM prepared plant materials; HK, TUI, JI, YHi, ET, MO, SU performed experiments and generated the data; HK, TUI, SU analyzed the data with help from TF, YS, MK, KY, SS, MN; HK, TUI, YH, SU, and YM wrote the paper; All authors discussed the results and commented on the manuscript.

## Figure legends

**Supplementary Fig. 1** | **Comparison of TKPR1 protein sequences from various plant species including angiosperms, gymnosperms, and phylogenetically basal plant species. a**, Alignment of TKPR1. All TKPR1 sequences had a conserved cysteine at the 82^nd^ residue of CjTKPR1 (red arrow), except for the *ms4* mutant. Eudicots and monocots (angiosperms) are highlighted in green and blue, respectively. Gymnosperms are highlighted in red. Asterisks indicate Japanese cedar. Black arrow and arrowhead indicate *Ceratopteris richardii* and *Marchantia polymorpha*, which are phylogenetically basal plant species that do not produce pollen. The putative NADP-binding domain is outlined. **b**, Amino acid identity and differences. Red boxes indicate comparison with CjTKPR1. Accession numbers are provided next to the species name. Multiple-sequence alignment was generated using CLC Main Workbench v22.0 software.

**Supplementary Fig. 2** | ***CjTKPR1* expression of complementation mutants in *A. thaliana***. The *ELONGATION FACTOR 1-alpha* (*AtEF1-α*) gene was used as an internal control (**a**). Expression in each *TKPR1* mutant was confirmed by RT-PCR (**b**) and RT-PCR-RFLP (**c**). CjWT, *CjTKPR1*-*WT*; CjWM, *CjTKPR1*-*WM*; CjMT, *CjTKPR1-MT*; CjMW, *CjTKPR1-MW* (Fig. 3a). Schematic structures of *AtEF1-α* (**d**) and *CjTKPR1* (**e**). Primer information is provided in Supplementary Table 5.

**Supplementary Table 1** | **Detailed information on the Japanese cedar materials used in this study**.

**Supplementary Table 2** | **Pollen production and single-nucleotide polymorphism (SNP) type for progenies of S8 (*ms4/ms4*) × S8HK5 (*Ms4/ms4*)**.

**Supplementary Table 3** | **Annotations of 67 genes in the *MS4* locus**.

**Supplementary Table 4** | **Pollen production and SNP types of selfing progenies of S8DY1(*Ms4/ms4*)**.

**Supplementary Table 5** | **Primers used in this study**.

**Supplementary Video 1** | **Time-lapse movie of flowering Japanese cedar male strobili**. Male strobili are initially closed (0–8 s). Then, the strobili begin to open, and microsporangia (yellow spherical structures) appear (8–14 s). Finally, microsporangia break open and pollen grains are released (14–23 s). The video was recorded using a TLC200 Pro camera (Brinno, Taipei, Taiwan) from 7^th^ to 10^th^, May 2020. Time points are indicated at the bottom of the video frame in the following format: yyyy/mm/dd, hh:mm:ss.

