## Supplementary Information for "Single-nucleotide substitution determines pollen production in Japanese cedar"

Kakui *et al.*

a

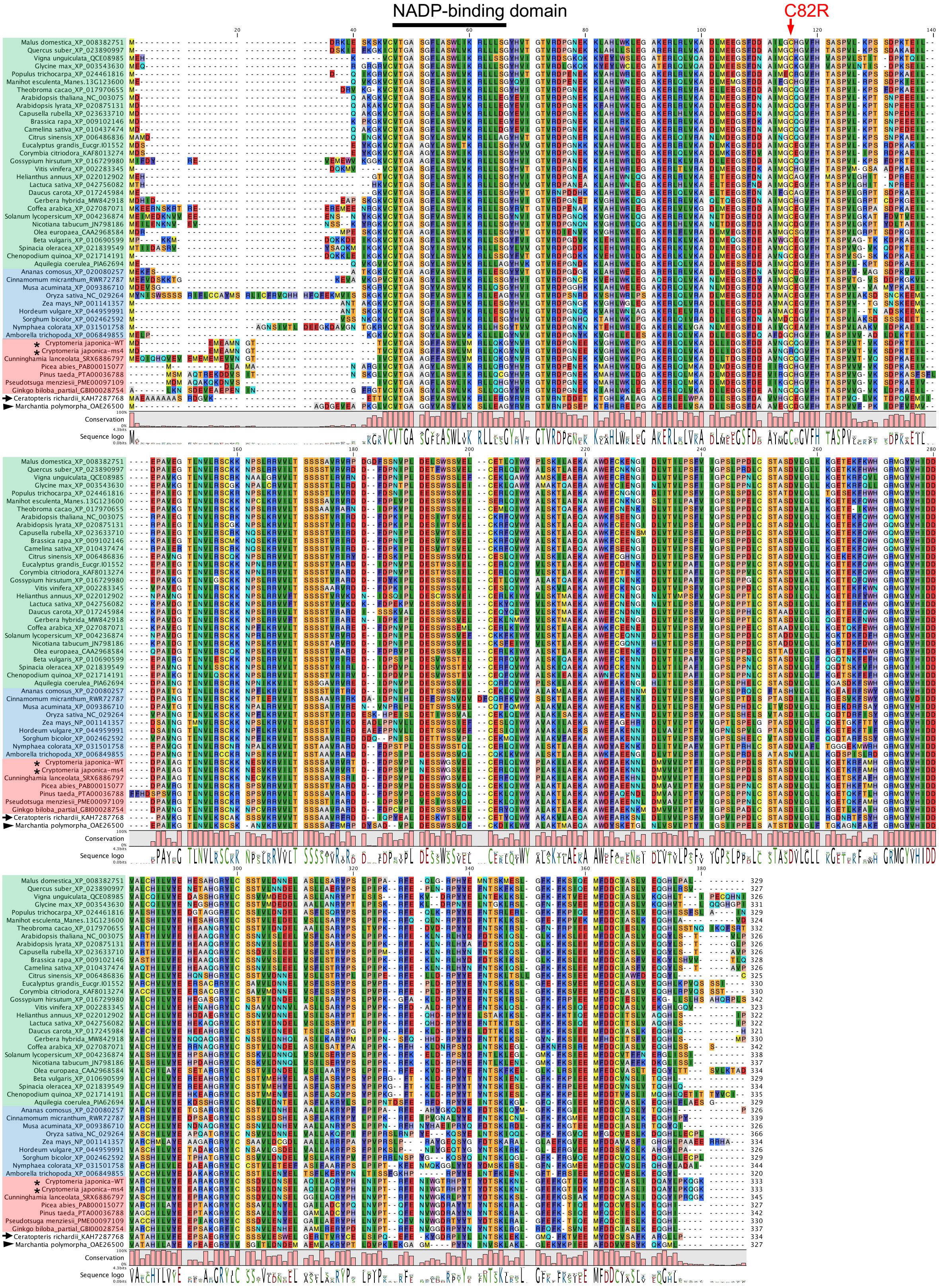

b

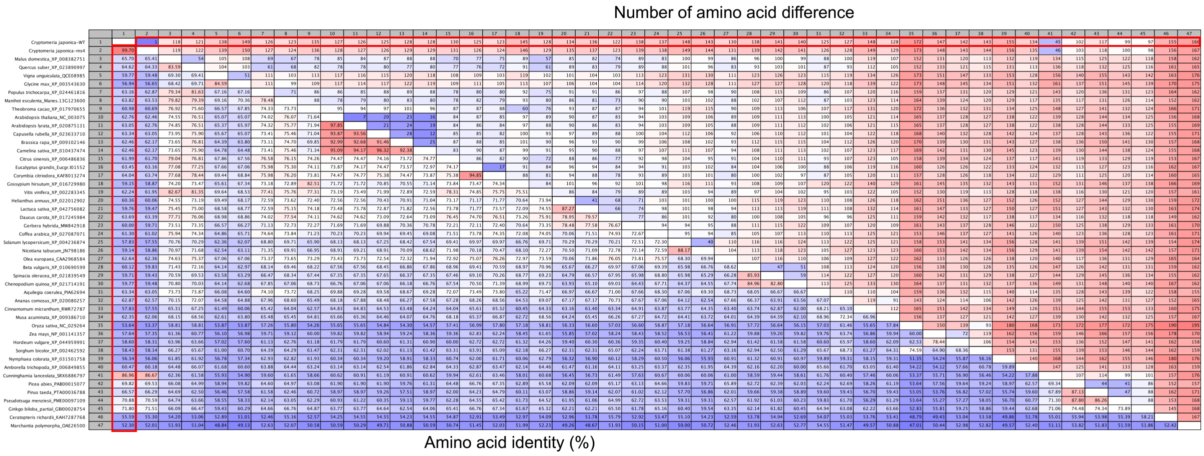


**Supplementary Fig. 1 | Comparison of TKPR1 protein sequences from various plant species including angiosperms, gymnosperms, and phylogenetically basal plant species. a**, Alignment of TKPR1. All TKPR1 sequences had a conserved cysteine at the 82$^{nd}$ residue of CjTKPR1 (red arrow), except for the *ms4* mutant. Eudicots and monocots (angiosperms) are highlighted in green and blue, respectively. Gymnosperms are highlighted in red. Asterisks indicate Japanese cedar. Black arrow and arrowhead indicate *Ceratopteris richardii* and *Marchantia polymorpha*, which are phylogenetically basal plant species that do not produce pollen. The putative NADP-binding domain is outlined. **b**, Amino acid identity and differences. Red boxes indicate comparison with CjTKPR1. Accession numbers are provided next to the species name. Multiple-sequence alignment was generated using CLC Main Workbench v22.0 software.

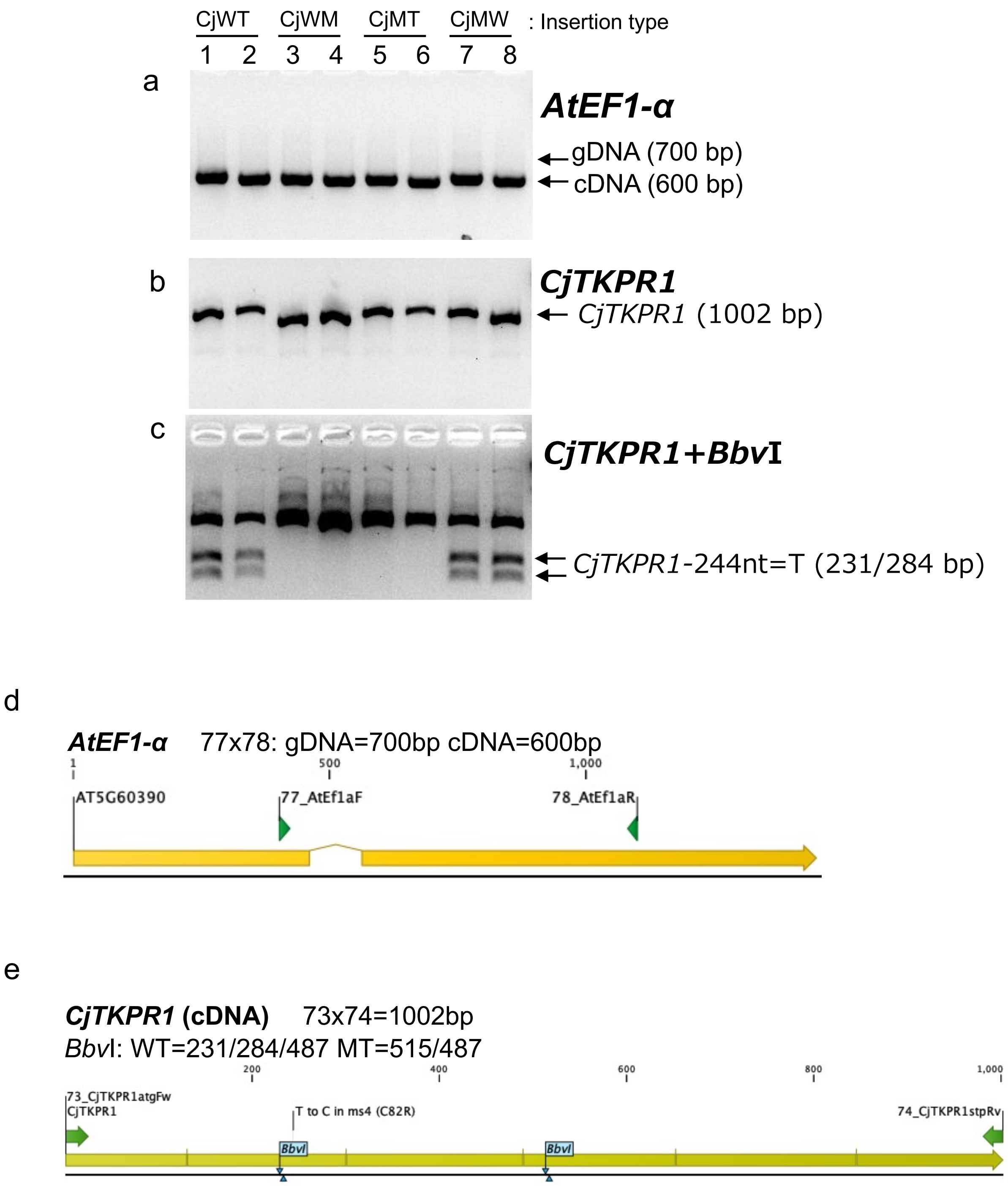


**Supplementary Fig. 2 | *CjTKPR1* expression of complementation mutants in *A.thaliana*.** The *ELONGATION FACTOR 1-alpha* (*AtEF1-α*) gene was used as an internal control (**a**). Expression in each *TKPR1* mutant was confirmed by RT-PCR (**b**) and RT-PCR-RFLP (**c**). CjWT, *CjTKPR1-WT*; CjWM, *CjTKPR1-WM*; CjMT, *CjTKPR1-MT*; CjMW, *CjTKPR1-MW* (Fig. 3a). Schematic structures of *AtEF1-α* (**d**) and *CjTKPR1* (**e**). Primer information is provided in Supplementary Table 5.

**Supplementary Table 1. Information of Japanese cedar materials in this study.**

| | Accession name | Abbreviation | Pollen production | SNP type of *CjTKPR1* (244-nt) | Purpose | Origin of accession |
|---|---|---|---|---|---|---|
| **Wild accessions** | Higashikanbara-5 | HK5 | Normal | T/T | Observation of pollen, counting of pollen number, Material for fine mapping | Niigata prefectural Forest Research Institute |
| | Dewanoyuki-1 | DY1 | Normal | T/T | Material for fine mapping | Yamagata prefectural Forest Research and Instruction Center |
| | Nakakubiki-4 | | Normal | T/T | Genotype and phenotype check | Niigata prefectural Forest Research Institute |
| | Santo-1 | | Normal | T/T | Histological analysis of male flower | Niigata prefectural Forest Research Institute |
| | Suzu-2 | | Normal | T/T | Genotype and phenotype check | Ishikawa Agriculture and Forestry Research Center Forestry Experiment Station |
| | Taisetsumurakami-4 | TM4 | Normal | T/T | Parent for RNA-seq | Niigata prefectural Forest Research Institute |
| | Shindai-8 (*ms4* mutant) | S8 | Pollen-free | C/C | *ms4* mutant, pollen observation, parent for fine mapping, linkage analysis, RNA-seq | Niigata University |
| | Shindai-3 (*ms1* mutant) | | Pollen-free | T/T | Genotype and phenotype check | Niigata University |
| | Shindai-1 (*ms2* mutant) | | Pollen-free | T/T | Genotype and phenotype check | Niigata University |
| | Shindai-5 (*ms3* mutant) | | Pollen-free | T/T | Genotype and phenotype check | Niigata University |
| **Crossing parents** | Shindai-8×Taisetsumurakami-4 | S8TM4 | Normal | T/C | Complementation test | Niigata prefectural Forest Research Institute |
| | Shindai-8×Higashikanbara-5 | S8HK5 | Normal | T/C | Parent for fine mapping | Niigata prefectural Forest Research Institute |
| | Shindai-8×Dewanoyuki-1 | S8DY1 | Normal | T/C | Parent for linkage analysis | Yamagata prefectual Forest Research and Instruction Center |
| **Crossing progenies** | P411 (progeny line of S8×S8HK5) | | Normal | T/C | Fine mapping, Electron microscopy | Forestry and Forest Products Research Institute |
| | P380 (progeny line of S8×S8HK5) | | Pollen-free | C/C | Fine mapping, Electron microscopy | Forestry and Forest Products Research Institute |

**Supplementary Table 2. Pollen production and SNP type for progenies of S8 (*ms4/ms4*) × S8HK5 (*Ms4/ms4*)**
Each individual data is provided as a Source Data file

| | SNP type at the 244-nt of *CjTKPR1* | |
|---|---|---|
| | T/C | C/C |
| Normal pollen | 46 | 0 |
| Pollen-free | 0 | 48 |

**Supplementary Table 3. Annotated sixty seven genes in the MS4 locus.**
*RNA-seq data were obtaind from Wei et al. 2021.

| | | | | | Male flower expression (TPM)* | | | Inner bark and leaf expression (TPM)* | | Homologous gene ID | | | | | |
|---|---|---|---|---|---|---|---|---|---|---|---|---|---|---|---|
| Gene No. | Transcript ID | Transcript strand(+/-) | Transcript length (bp) | Distance from AX-174206035 (bp) | Nakakubiki-4 (2011/10/4) | Nakakubiki-4 (2011/10/13) | Nakakubiki-4 (2011/10/19) | Ooi-7 (2016/6/24) | S8HK5 (2016/6/20) | *Picea abies* | *Pinus taeda* | *Oryza sativa* | *Arabidopsis thaliana* | Gene name (*A.thaliana*) | Annotation (*A.thaliana*) |
| 1 | CJt019505 | + | 885 | 277,748 | 35.69 | 11.54 | 9.90 | 53.11 | 55.40 | PAB00035884 | ND | ND | AT5G02640 | - | hypothetical protein |
| 2 | CJt083246 | - | 837 | 420,432 | 3.39 | 1.11 | 1.01 | 4.14 | 11.19 | ND | ND | ND | AT5G02640 | - | hypothetical protein |
| 3 | CJt104934 | + | 397 | 685,277 | 7.90 | 3.02 | 0.27 | 2.97 | 6.23 | ND | PTA00035456 | ND | ND | | |
| 4 | CJt022182 | - | 1461 | 808,783 | 2.09 | 1.68 | 1.86 | 2.10 | 2.96 | ND | ND | ND | ND | | |
| 5 | CJt086258 | - | 848 | 809,906 | 0.00 | 0.00 | 0.00 | 0.00 | 0.00 | ND | ND | ND | ND | | |
| 6 | CJt116102 | + | 1390 | 810,311 | 0.00 | 0.00 | 0.00 | 0.00 | 0.22 | ND | ND | ND | ND | | |
| 7 | CJt108958 | + | 1672 | 810,334 | 0.56 | 0.48 | 0.26 | 1.90 | 1.34 | ND | ND | ND | ND | | |
| 8 | CJt113461 | + | 1608 | 810,378 | 1.44 | 1.40 | 0.79 | 1.30 | 1.51 | ND | ND | ND | ND | | |
| 9 | CJt103230 | - | 1697 | 813,945 | 25.30 | 14.99 | 17.36 | 13.03 | 17.57 | PAB00062857 | ND | ND | AT1G10150 | - | Carbohydrate-binding protein |
| 10 | CJt098789 | - | 1741 | 814,061 | 3.27 | 2.04 | 2.78 | 3.98 | 4.22 | PAB00064377 | ND | ND | AT1G10150 | - | Carbohydrate-binding protein |
| 11 | CJt075881 | - | 294 | 880,460 | 21.92 | 14.03 | 7.12 | 4.90 | 13.94 | ND | ND | ND | ND | | |
| 12 | CJt069068 | - | 328 | 1,037,514 | 0.00 | 0.34 | 0.26 | 0.18 | 0.00 | ND | ND | ND | ND | | |
| 13 | CJt100969 | - | 480 | 1,068,450 | 0.00 | 0.00 | 0.20 | 0.43 | 1.24 | ND | ND | ND | ND | | |
| 14 | CJt065237 | - | 315 | 1,198,817 | 0.35 | 0.00 | 0.39 | 1.08 | 0.19 | ND | ND | ND | ND | | |
| 15 | CJt108163 | - | 7100 | 1,207,780 | 0.28 | 0.29 | 0.16 | 0.27 | 3.30 | ND | ND | ND | ATMG00860 | ORF158 | DNA/RNA polymerases superfamily protein |
| 16 | CJt000893 | - | 677 | 1,317,579 | 0.45 | 0.12 | 0.20 | 0.90 | 1.55 | ND | ND | ND | ATMG00750 | ORF119 | GAG/POL/ENV polyprotein |
| 17 | CJt114371 | + | 617 | 1,322,053 | 0.20 | 0.63 | 0.47 | 1.57 | 1.36 | ND | ND | ND | ND | | |
| 18 | CJt001465 | + | 895 | 1,368,671 | 0.84 | 0.58 | 0.92 | 1.62 | 1.77 | ND | ND | ND | ND | | |
| 19 | CJt041627 | + | 329 | 1,368,699 | 0.44 | 1.14 | 0.00 | 1.31 | 1.82 | ND | ND | ND | ND | | |
| 20 | CJt046182 | - | 308 | 1,401,470 | 0.00 | 0.00 | 0.00 | 0.64 | 0.96 | ND | ND | ND | ND | | |
| 21 | CJt082717 | - | 897 | 1,418,359 | 0.22 | 0.00 | 0.25 | 0.41 | 0.48 | ND | ND | ND | ND | | |
| 22 | CJt013759 | - | 4093 | 1,426,583 | 7.71 | 6.01 | 5.01 | 30.27 | 25.78 | PAB00028684 | PTA00075208 | ND | AT5G58140 | PHOT2 | Membrane-bound protein serine/threonine kinase that functions as blue light photoreceptor in redundancy with PHO1. |
| 23 | CJt084897 | + | 1553 | 1,711,794 | 44.86 | 51.75 | 44.61 | 56.45 | 51.12 | PAB00000046 | ND | OS07G38620 | AT3G03890 | HOZ | Dimeric β-barrel protein that is structurally related to the putative non-canonical heme oxygenase (HO) and is located in chloroplasts. |
| 24 | CJt053658 | - | 937 | 1,727,608 | 0.81 | 0.83 | 0.42 | 0.75 | 0.83 | ND | ND | ND | AT1G67020 | - | transmembrane protein |
| 25 | CJt050936 | - | 377 | 1,868,970 | 0.26 | 0.27 | 0.58 | 0.46 | 0.55 | ND | ND | ND | ND | | |
| 26 | CJt014186 | + | 921 | 1,898,628 | 0.08 | 0.00 | 0.71 | 0.48 | 0.42 | ND | ND | ND | ND | | |
| 27 | CJt056754 | + | 1217 | 1,944,641 | 0.58 | 1.11 | 0.87 | 0.94 | 1.03 | ND | ND | ND | AT1G67020 | - | transmembrane protein |
| 28 | CJt010501 | + | 384 | 1,948,218 | 0.75 | 0.21 | 0.19 | 0.54 | 1.55 | ND | ND | ND | ND | | |
| 29 | CJt085228 | - | 539 | 2,010,758 | 7.13 | 5.11 | 2.74 | 1.35 | 2.42 | ND | ND | ND | AT2G23755 | - | transmembrane family 220 helix protein |
| 30 | CJt106991u | + | 699 | 2,055,095 | 3.31 | 2.81 | 2.18 | 4.10 | 3.18 | ND | ND | ND | ND | | |
| 31 | CJt037660u | + | 505 | 2,055,731 | 0.00 | 0.30 | 0.04 | 0.00 | 0.27 | ND | ND | ND | ND | | |
| 32 | CJt074706 | - | 627 | 2,111,177 | 0.34 | 0.00 | 0.00 | 0.51 | 1.22 | ND | ND | ND | ND | | |
| 33 | CJt068182 | - | 484 | 2,111,231 | 1.01 | 0.60 | 0.23 | 1.49 | 0.97 | ND | ND | ND | ND | | |
| 34 | CJt107062 | + | 427 | 2,157,527 | 0.00 | 0.00 | 0.00 | 0.51 | 0.50 | ND | ND | ND | ND | | |
| 35 | CJt072915 | - | 341 | 2,168,439 | 1.53 | 0.32 | 0.69 | 0.95 | 2.76 | ND | ND | ND | AT3G22220 | - | hAT transposon superfamily |
| 36 | CJt009333 | + | 379 | 2,183,228 | 0.77 | 0.27 | 0.29 | 2.05 | 2.60 | ND | ND | ND | ND | | |
| 37 | CJt022014 | + | 687 | 2,188,638 | 0.00 | 0.00 | 0.58 | 1.13 | 1.23 | ND | ND | ND | ND | | |
| 38 | CJt043927 | - | 344 | 2,215,844 | 15.60 | 17.09 | 19.96 | 13.32 | 14.14 | ND | ND | ND | ND | | |
| 39 | CJt096053 | - | 368 | 2,504,649 | 0.00 | 0.28 | 0.30 | 0.00 | 7.56 | ND | ND | ND | ND | | |
| 40 | CJt087566 | - | 1624 | 2,600,989 | 22.33 | 28.07 | 24.49 | 52.02 | 54.03 | PAB00008938 | PTA00029192 | OS10G01044 | AT3G18890 | Tic62 | NAD(P)-binding Rossmann-fold superfamily protein |
| 41 | CJt001736 | - | 580 | 2,610,405 | 0.36 | 0.16 | 0.26 | 1.51 | 1.19 | ND | ND | ND | AT4G23160 | CRK8 | Encodes a cysteine-rich receptor-like protein kinase. |
| 42 | CJt009105 | - | 330 | 2,610,480 | 0.00 | 0.00 | 0.00 | 0.00 | 0.00 | ND | ND | ND | AT4G23160 | CRK8 | Encodes a cysteine-rich receptor-like protein kinase. |
| 43 | CJt087973 | - | 2221 | 2,791,991 | 19.04 | 14.55 | 18.34 | 24.63 | 51.02 | PAB00028559 | ND | OS03G55950 | AT1G63850 | - | BTB/POZ domain-containing protein |
| 44 | CJt020290 | - | 312 | 2,797,453 | 0.00 | 0.00 | 0.00 | 0.16 | 0.46 | ND | ND | ND | ND | | |
| 45 | CJt000212 | + | 1421 | 3,164,822 | 14.06 | 5.74 | 8.63 | 34.79 | 25.06 | ND | ND | ND | AT1G63840 | - | RING/U-box superfamily protein |
| 46 | CJt031649 | + | 331 | 3,318,747 | 1.28 | 1.34 | 1.43 | 0.42 | 0.00 | ND | ND | ND | ND | | |
| 47 | CJt086070 | - | 4774 | 3,347,851 | 23.99 | 23.51 | 19.15 | 32.07 | 26.83 | PAB00016736 | PTA00007814 | OS04G57140 | AT5G65930 | ZWI | encodes a novel member of the kinesin superfamily of motor proteins. recessive mutations have reduced number of trichome branches. |
| 48 | CJt060254 | + | 298 | 4,003,457 | 2.68 | 1.61 | 1.71 | 1.17 | 1.01 | ND | ND | ND | ND | | |
| 49 | CJt002142 | - | 775 | 4,014,073 | 0.17 | 0.50 | 0.80 | 1.77 | 2.44 | ND | ND | ND | ND | | |
| 50 | CJt002157 | + | 6433 | 4,250,099 | 5.84 | 4.82 | 3.62 | 8.59 | 4.67 | PAB00013806 | PTA00009996 | OS01G67970 | AT3G48430 | REF6 | Relative of Early Flowering 6 (REF6) encodes a Jumonji N/C and zinc finger domain-containing protein that acts as a positive regulator of flowering in an FLC-dependent pathway. |
| 51 | CJt037308 | + | 377 | 4,791,393 | 0.00 | 1.09 | 0.00 | 0.00 | 0.00 | ND | ND | ND | ND | | |
| 52 | CJt034360 | + | 1169 | 4,800,363 | 338.71 | 1131.88 | 183.17 | 0.54 | 0.00 | PAB00015077 | PTA00036788 | OS08G40440 | AT4G35420 | TKPR1 | Involved in the biosynthesis of hydroxylated tetraketide compounds that serve as sporopollenin precursors (the main constituents of exine).<br>Is essential for pollen wall development. Acts on tetraketide alpha-pyrones and reduces the carbonyl function on the tetraketide alkyl chain to a secondary alcohol function. |
| 53 | CJt114344 | - | 503 | 4,806,485 | 8.31 | 6.72 | 3.64 | 2.11 | 2.98 | ND | ND | ND | ND | | |
| 54 | CJt053498 | - | 492 | 4,807,761 | 0.17 | 0.91 | 0.20 | 0.23 | 0.55 | ND | ND | ND | AT3G26630 | - | Tetratricopeptide repeat (TPR)-like superfamily protein |
| 55 | CJt098949 | - | 665 | 4,809,658 | 0.23 | 1.08 | 0.13 | 0.15 | 0.30 | ND | ND | ND | AT1G71420 | - | Tetratricopeptide repeat (TPR)-like superfamily protein |
| 56 | CJt028952 | + | 381 | 4,811,348 | 0.77 | 1.35 | 0.29 | 0.23 | 0.54 | ND | ND | ND | ND | | |
| 57 | CJt097191 | + | 313 | 4,811,634 | 0.35 | 0.00 | 0.00 | 0.16 | 0.94 | ND | ND | ND | ND | | |
| 58 | CJt035612 | + | 579 | 4,987,752 | 2.46 | 6.64 | 4.81 | 0.00 | 0.14 | ND | ND | ND | ND | | |
| 59 | CJt102965 | + | 363 | 4,987,773 | 3.18 | 4.73 | 2.34 | 1.38 | 2.04 | ND | ND | ND | ND | | |
| 60 | CJt031343 | + | 352 | 5,163,119 | 0.58 | 1.19 | 0.65 | 0.15 | 0.62 | ND | ND | ND | AT3G01410 | - | Polynucleotidyl transferase, ribonuclease H-like superfamily protein |
| 61 | CJt079034 | - | 1370 | 5,288,454 | 4.82 | 3.37 | 4.12 | 4.48 | 7.15 | PAB00015943 | PTA00044917 | OS09G32025 | AT2G26540 | HEMD | Encodes a uroporphyrinogen-III synthase involved in tetrapyrrole biosynthesis. The protein localizes to the chloroplast. Member of the plant-specific DUF724 protein family. |
| 62 | CJt041442 | - | 589 | 5,441,202 | 0.50 | 0.40 | 0.30 | 0.38 | 0.53 | ND | ND | ND | ND | | |
| 63 | CJt106494 | - | 431 | 5,452,776 | 0.42 | 0.22 | 0.47 | 0.65 | 2.35 | ND | ND | ND | AT3G57120 | - | Protein kinase superfamily protein |
| 64 | CJt107266 | - | 1895 | 5,453,169 | 0.87 | 0.53 | 0.19 | 0.53 | 3.13 | ND | PTA00039094 | OS02G57440 | AT1G51940 | LYK3 | Encodes a LysM-containing receptor-like kinase. Induction of chitin-responsive genes by chitin treatment is not blocked in the mutant. |
| 65 | CJt068591 | - | 329 | 5,484,010 | 1.13 | 0.34 | 0.00 | 0.93 | 1.05 | ND | PTA00004023 | ND | ND | | |
| 66 | CJt085666 | + | 3114 | 7,648,181 | 17.31 | 9.12 | 11.07 | 38.08 | 39.21 | PAB00035566 | PTA00036585 | OS03G52239 | AT2G23760 | BLH4 | Encodes a member of the BEL family of homeodomain proteins. Plants doubly mutant for saw1/saw2 (blh2/blh4) have serrated leaves. |
| 67 | CJt113594 | + | 698 | 7,650,117 | 0.21 | 0.11 | 0.00 | 0.11 | 1.24 | ND | ND | ND | ND | | |

**Supplementary Table 4. Pollen production and SNP type for selfing progenies of S8DY1(*Ms4/ms4*)**
Each individual data is provided as a Source Data file

| | SNP type at the 244-nt of *CjTKPR1* | | |
|---|---|---|---|
| | T/T | T/C | C/C |
| Normal pollen | 4 | 13 | 0 |
| Pollen-free | 0 | 0 | 3 |

**Supplementary Table 5**. Primer list in this study.

| Target | Purpose | Template | Forward primer | | Reverse primer | |
|---|---|---|---|---|---|---|
| | | | Primer name | sequence (5'-3') | Primer name | sequence (5'-3') |
| *CjTKPR1* | RT-PCR | cDNA/Japanese cedar/various tissue | 143_CjTKPR1f | ACTGAAATTCTGGATCCAGCTATTGC | 144_CjTKPR1r | CTTTTAACAACCCAAGCACATC |
| *CjEF1-alpha* | RT-PCR (internal control) | cDNA/Japanese cedar/various tissue | 145_CjEf1a-f | CAGAGAGCATGCTCTTCTGGCC | 146_CjEf1a-r | CATTGAACCCCACATTGTCACCAGGC |
| *CjTKPR1* | *in situ* hybridization probe | cDNA/ Japanese cedar | CjTKPR1_probe w/SP6-Fw | ATTTAGGTGACACTATAGAACATATGGGGAACAAGG | CjTKPR1_probe w/T7-Rev | TAATACGACTCACTATAGGGTTATTTTCCTTGTTTAGGAAG |
| *AtTKPR1* promoter | complementation/vector construction | gDNA/*Arabidopsis thaliana* /leaf | 12_AtTKPR1proIFfw-1 | agtcgactggatccgGTCAAGTTCCTCTTACAGCAGTAC | 9_AtTKPR1proR1 | TTTCCGGTATAAATGGAATCACACCCG |
| *CjTKPR1* coding | complementation/vector construction | cDNA/Japanese cedar(WT or Shindai-8)/Flower buds | 15_CjTKPR1atgFwIF | CATTTATACCGGAAAATGGATGAAATGGAAGCCATGAATG | 16_CjTKPR1stpRvIF | AATGTTTGAACGATCTTATTTTCCTTGTTTAGGAAGG |
| *CjTKPR1* coding (T244C) | complementation/vector construction | Plasmid/pENTR:AtTKPR1pro:CjTKPR1coding(WT):NosT | 24_CjTKPR1rv | ACAGCATCATCAAAGCTACC | 25_CjTKPR1TtoCf | CAATGGCcGCCAGGGAGTTTTTCACAC |
| *CjTKPR1* coding (C244T) | complementation/vector construction | Plasmid/pENTR:AtTKPR1pro:CjTKPR1coding(Shindai-8):NosT | 59_CjTKPR1mtrv2true | ACAGCgTCATCAAAGCTACC | 26_CjTKPR1CtoT4mtFw | CAATGGCTGCCAGGGAGTTTTTCACAC |
| Nos teminator | complementation/vector construction | pGWB3 | 17_NosTFw | GATCGTTCAAACATTTGGC | 18_NosTrPluspENTR-IF | GTCTAGATATCTCGAGATCTAGTAACATAGATGAC |
| *AtTKPR1* wild-type | genotyping for AtTKPR1 wild-type | gDNA/*Arabidospsis thaliana* | 27_AtTKPR1f_CL194 | GAAGAAACTTGCGCACCTATG | 60_AtTKPR1r | TGGACCCAAAAACGAGTCAT |
| *AtTKPR1 tkpr1-1* | genotyping for *tkpr1-1* | gDNA/*Arabidospsis thaliana* | 62_LB1forTKPR1r | GCCTTTTCAGAAATGGATAAATAGCCTTGCTTCC | 60_AtTKPR1r | TGGACCCAAAAACGAGTCAT |
| AtTKPR1 | genotyping for external TKPR1(exon-junction primer) | gDNA with transgene/*Arabidopsis thaliana* | 69_AtTKPR1JF1 | ACAGTCAGAGATCCAGGAAATG | 70_AtTKPR1JR1 | TATTTAGTTTCTCAAACCTCTTGG |
| *AtEF1-alpha* | Expression control of complementation analysis | cDNA/ *Arabidopsis thaliana* | 77_AtEf1aF | TGAGCACGCTCTTCTTGCTTTCA | 78_AtEf1aR | ATGATGACCTGGGAGGTGAAG |
| *CjTKPR1* | Expression check of comlementation test | cDNA/*Arabidopsis thaliana* (with *CjTKPR1* ) | 73_CjTKPR1atgFw | ATGGATGAAATGGAAGCCATGAATG | 74_CjTKPR1stpRv | TTATTTTCCTTGTTTAGGAAGG |
| CJt085666 | Linkage mapping | gDNA/ Japanese cedar | CJt085666_F | ATGTCACAACAGACGAGAAGGTT | CJt085666_R | CGCAAATTGGTGGTTATTATTGT |
